# Multiomic single-cell analysis identifies von Willebrand factor and TIM3-expressing *BCR-ABL1*^+^ CML stem cells

**DOI:** 10.1101/2023.09.14.557507

**Authors:** Malin S. Nilsson, Hana Komic, Johan Gustafsson, Zahra Sheybani, Sanchari Paul, Ola Rolfson, Kristoffer Hellstrand, Lovisa Wennström, Anna Martner, Fredrik B. Thorén

**Affiliations:** TIMM Laboratory at Sahlgrenska Center for Cancer Research, University of Gothenburg, Gothenburg, Sweden; Department of Microbiology and Immunology, Institute of Biomedicine, Sahlgrenska Academy, University of Gothenburg, Gothenburg, Sweden; Department of Medical Biochemistry and Cell Biology, Institute of Biomedicine, Sahlgrenska Academy, University of Gothenburg, Gothenburg, Sweden; Department of Biology and Biological Engineering, Chalmers University of Technology, Gothenburg, Sweden; Department of Orthopaedics, Institute of Clinical Sciences, Sahlgrenska Academy, University of Gothenburg, Gothenburg, Sweden; Department of Infectious Diseases, Institute of Biomedicine, Sahlgrenska Academy, University of Gothenburg, Gothenburg, Sweden; Department of Hematology, Sahlgrenska University Hospital, Gothenburg, Sweden

**Keywords:** CML, BCR-ABL1, CITE-seq, multiomic, LSC, von Willebrand factor, TIM3

## Abstract

Tyrosine kinase inhibitors (TKI) only rarely eradicate leukemic stem cells (LSC) in chronic myeloid leukemia (CML) which commonly necessitates life-long therapy and monitoring of patients. Understanding details of leukemic hematopoiesis in CML may identify targetable pathways for sustained LSC elimination. This study utilized multiomic single-cell characterization of the CD14^-^CD34^+^ hematopoietic stem and progenitor cell (HSPC) compartment in CML. Combined proteo-transcriptomic profiling of 597 genes and 51 proteins (CITE-seq) was performed along with parallel detection of *BCR-ABL1* transcripts in 70,000 HSPC from 16 chronic phase patients and five healthy controls. CD14^-^CD34^+^ HSPC from diagnosis samples displayed distinct myeloid cell bias with cells mainly annotated as LSC, lympho-myeloid progenitors (LMP)-II, erythrocyte and megakaryocyte progenitors, while few hematopoietic stem cells (HSC), LMP-I, dendritic cell or B cell progenitors were detected. In-depth analysis of the immature CD14^-^CD34^+^CD38^-/low^ compartment revealed two distinct populations of *BCR-ABL1*-expressing CML LSC (denoted LSC-I and LSC-II), where LSC-I showed features of quiescence and CD45RA^-^cKIT^-^CD26^+^ TKI therapy-resistant phenotype. These subtypes of immature LSC showed high surface expression of TIM3 and transcription of the von Willebrand factor gene (*VWF*). Our findings imply that expression of *VWF* and TIM3 distinguish LSC from HSC and may be linked to aberrant myeloid-biased hematopoiesis in CML. Additionally, the results identify TIM3 as a conceivable target for sustained elimination of immature LSC in CML.

**Key points:**

- We present a method to detect *BCR-ABL1* expression at the single-cell level that is compatible with high-throughput CITE-seq
- The most immature *BCR-ABL1*-expressing LSC population in primary CML shows enhanced expression of von Willebrand factor and TIM3

## Introduction

Chronic myeloid leukemia (CML) is caused by chromosomal translocation and molecular juxtapositioning of *ABL1* on chromosome 9 and *BCR* on chromosome 22^1^. The resulting *BCR-ABL1* fusion oncogene yields constitutive activation of the ABL1 tyrosine kinase that promotes proliferation and survival of leukemic cells^2^. CML is believed to be initiated in immature hematopoietic stem cells (HSC) resulting in progressive overgrowth of mature granulocytic cells and their precursors^3,4^.

The long-term survival in CML has dramatically improved since the introduction of BCR-ABL1-targeted tyrosine kinase inhibitors (TKI)^5^, but CD34^+^CD38^-^ TKI-resistant and quiescent leukemic stem cells (LSC) mostly persist despite TKI therapy^3^. As remaining LSC may rekindle disease upon treatment discontinuation, CML patients typically require life-long TKI therapy^6^ and clinical monitoring. Several studies have thus focused on defining unique features of LSC that may allow their targeted elimination, potentially curing the disease. Analysis of LSC is challenging as they constitute a minor fraction of leukemic cells and are difficult to distinguish from normal HSC. However, advances in single-cell technologies have enabled the identification of gene and cell surface marker signatures of CML stem cells that associate with resistance to TKI therapy^7–9^, which may translate into the identification of novel therapeutic targets.

A further challenge for in-depth studies of CML LSC is that the disease-causing translocation is found approximately 5 kb from the poly(A) tail of the *BCR-ABL1* transcript. As most contemporary high-throughput single-cell RNA sequencing (scRNA-seq) methods rely on amplification and sequencing of the 3’ end of transcripts, characterization of leukemic cells by parallel detection of the leukemic translocation is problematic. To overcome this challenge, we adapted the ‘Genotyping of Transcriptomes’ (GoT) method^10^ to combine *BCR-ABL1* expression detection at the single-cell level with CITE-seq (cellular indexing of transcriptomes and epitopes by sequencing) of >70,000 hematopoietic stem and progenitor cells (HSPC) from CML patients and healthy controls. We report that CML hematopoiesis occurred via LSC and myeloid-biased lympho-myeloid progenitors (LMP)-II towards myeloid, megakaryocyte and erythrocyte progenitors. High surface expression of TIM3 and transcription of the von Willebrand Factor gene (*VWF*) defined immature populations of LSC and may explain the observed bias towards myeloid differentiation. Our results may point towards conceivably targetable characteristics of immature CML stem cells.

## Methods

### Primary cells and cell lines

Chronic phase bone marrow (BM) samples were aspirated from the posterior iliac crest at CML diagnosis (n=16) and 3-7 months into TKI treatment (n=12) at the Sahlgrenska University Hospital (Gothenburg, Sweden) and Uddevalla Hospital (Uddevalla, Sweden). Healthy BM samples were obtained from the femur of osteoarthritic patients (n=5) undergoing hip replacement surgery at the Sahlgrenska University Hospital (Mölndal, Sweden). The study was approved by the Regional Ethics Committee in Gothenburg and all participants gave written informed consent. Clinical details of patients and healthy donors are provided in Supplemental Table 1. BM mononuclear cells (MNC) were isolated by density gradient centrifugation followed by CD15^+^ granulocyte removal and/or enrichment of CD34^+^ cells before cryopreservation. Cryopreserved aliquots of the CML cell line K562 were used as controls.

### Library preparation for single-cell CITE-seq analysis

Following isolation of CD14^-^CD34^+^ HSPC by MACS technology (Miltenyi Biotec) and FACS, the BD Rhapsody Single-Cell Analysis System (BD Biosciences) was used for multiomic CITE-seq analysis of 597 genes and 51 surface proteins (Figure 1A). The panels (Supplemental Tables 2-4) comprised lineage markers and previously reported LSC biomarkers as well as markers related to stemness, proliferation, and redox status^11–13^. Supplemental Table 5 lists antibodies used for FACS-sorting.

**Figure 1.**
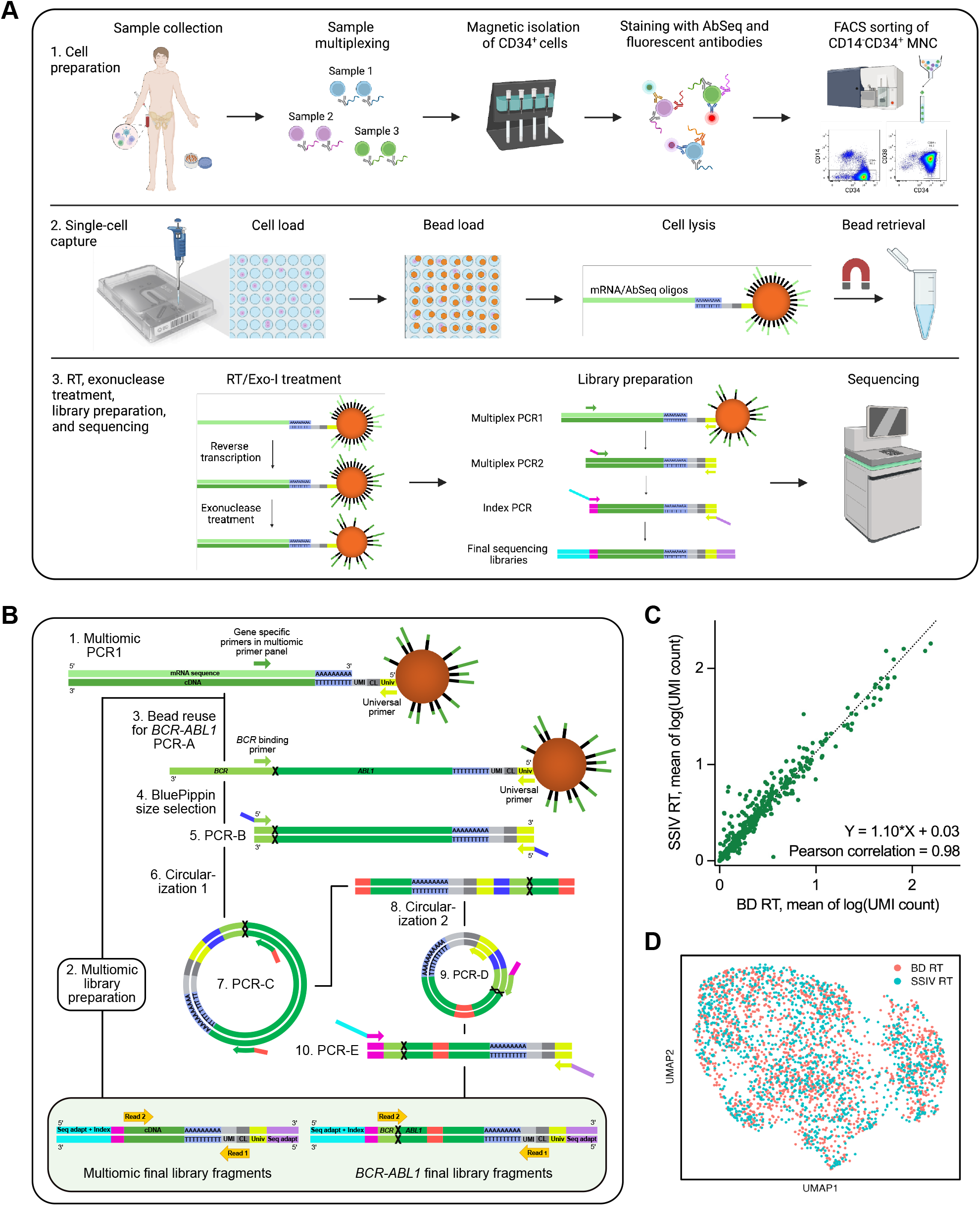
Multiomic CITE-seq analysis paired with *BCR-ABL1* detection at the single-cell level. (A) Overview of the workflow employed for CITE-seq analysis of CD14^-^CD34^+^ HSPC from healthy and CML patient BM. Created with BioRender.com. (B) Illustration of the method set up and employed for combined *BCR-ABL1* detection and CITE-seq. UMI, unique molecular identifier; CL, cell label; Univ, universal oligo. Sequences are not proportional in length. (C) Correlation of expression data for the 597 genes in the multiomic mRNA panel generated from K562 cells using the original BD Rhapsody reverse transcription procedure (BD RT; n=1,716) or SuperScript IV RT (SSIV RT; n=1,363). (D) Unsupervised clustering and UMAP visualization of K562 mRNA expression data generated using BD or SSIV RT.

### BCR-ABL1 sequencing library preparation

To assess *BCR-ABL1* expression in parallel with CITE-seq, we developed a method inspired by that reported by Nam *et al*.^10^, using PCR and circularization reactions to transpose the *BCR-ABL1* fusion point closer to the 3’ end (Figure 1B; with details provided in Supplemental Methods). Supplemental Table 6 provides the sequences of primers used in the workflow.

### Multiomic data analysis

Multiomic sequencing data was processed using the BD Rhapsody Targeted Analysis Pipeline (BD Biosciences, v. 1.10.1). After filtering cells based on genes expressed vs library size in SeqGeq software (BD Biosciences, v. 1.8.0)(Supplemental Figure 1), data analysis was performed in Rstudio (v. 2022.07.1+554; R v. 4.2.1) using the Seurat package (v. 4.2.0)^14^ as described in the Supplemental Methods.

### Statistical analysis

Statistical analyses were performed in Rstudio and GraphPad Prism (v. 9.5.1). Differentially expressed genes and proteins were assessed by the non-parametric Wilcoxon Rank Sum test in R using Seurat’s FindAllMarkers and FindMarkers functions. Differences in cell type proportions were assessed in GraphPad Prism using the unpaired Mann-Whitney test.

## Results

### Detection of BCR-ABL1 transcripts in single cells with paired multiomic expression data

Decisive distinction of CML-derived from healthy stem cells within the leukemic BM requires assessment of *BCR-ABL1* expression at the single-cell level. We performed 3’ end capture-based multiomic CITE-seq analysis of 597 genes and 51 proteins in multiplexed CML and healthy BM samples (Figure 1A). However, this methodology typically only provides sequence information close to transcript ends, preventing the detection of the *BCR-ABL1* transcript-specific fusion point located >5 kb from the poly(A) tail.

In order to simultaneously detect *BCR-ABL1* transcripts in single cells subjected to CITE-seq, we used a series of PCR and circularization reactions to move the fusion point closer to the cell capture bead cell label and enable *BCR-ABL1*-specific short-read sequencing (Figure 1B). Cell labels from the separate datasets enabled coupling of *BCR-ABL1* expression to the CITE-seq-derived targeted gene and protein expression data.

To enable long-range reverse transcription (RT), we replaced the reverse transcriptase of the original workflow with a high-processivity transcriptase (SuperScript IV; SSIV), which allowed efficient amplification of the ∼5,300 bp product required for the first *BCR-ABL1*-specific PCR (PCR-A; Figure 1B). The alternative RT procedure did not affect the performance of the multiomic analysis, as sequencing libraries obtained from CML-derived K562 cells using the original and alternative RT procedure showed satisfactory correlation between expression of the 597 genes in the targeted mRNA panel (Pearson correlation = 0.98; Figure 1C), and there were no signs of separate clustering in dimensionality reduction analysis (Figure 1D). Interestingly, linear regression analysis of the correlation plot indicated overall increased sensitivity with the adapted RT procedure (Figure 1C), additionally illustrated by the proportion of *ABL1*^+^ K562 being strikingly higher using the alternative RT procedure (92% vs 69.5%).

On four cartridges, spike-ins of K562 cells were included to assess *BCR-ABL1* detection sensitivity. *BCR-ABL1* transcripts were detected in 19.8, 40.8 and 49.8% of K562 cells on three cartridges, while *BCR-ABL1* transcript detection sensitivity was low on the fourth cartridge (0.97% positive K562 cells). In healthy donor BM cells, 0.24% of cells showed *BCR-ABL1* expression, likely reflecting some false positivity.

### Generation of healthy and CML-specific annotation references

CITE-seq and *BCR-ABL1* expression data were generated for CD14^-^CD34^+^ HSPC obtained from BM samples from CML patients at diagnosis (n=16; 58,682 cells after quality filtering) and during TKI treatment (n=12; 4,571 cells) as well as from healthy controls (n=5; 7,161 cells).

To enable subsequent rough annotation of leukemic cells, we first created a healthy annotation reference using data from the five healthy BM samples (Figure 2A). Cluster annotations were determined based on previous publications as described in the Supplemental Methods, with annotation marker dot plots, feature plots, cell cycle analysis and cluster differential gene and protein expression data shown in Figure 2B-C, Supplemental Figure 2A-C and Supplemental Tables 7-8. Despite only using mRNA expression data for clustering, also AbSeq-derived protein expression appeared confined to cell-specific portions of the UMAP (Supplemental Figure 2C), supporting the validity of the clustering and analysis. The cell annotations in Figure 2A are henceforth referred to as the ‘healthy reference’.

**Figure 2.**
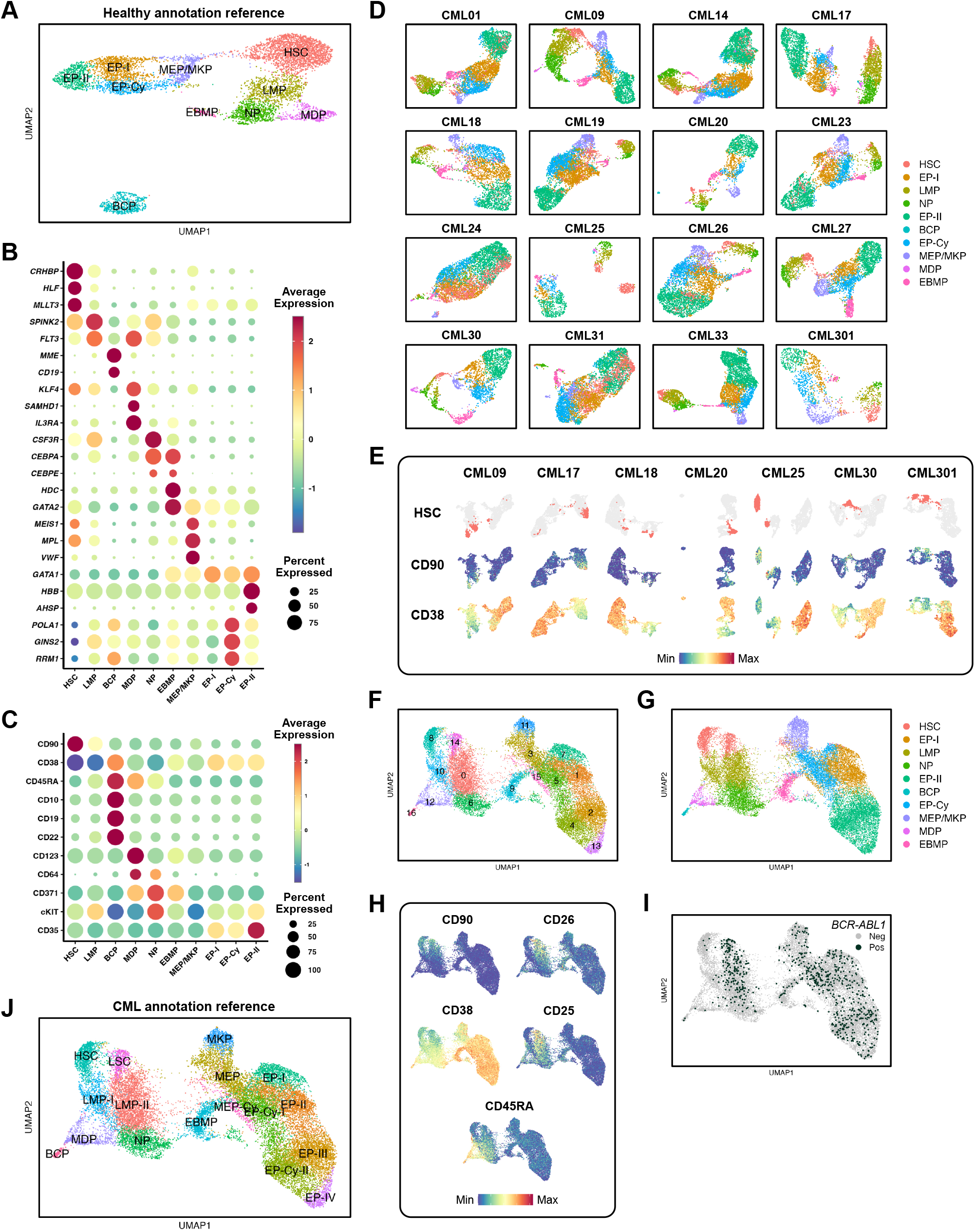
Generation of healthy and CML-specific annotation references. (A-C) Generation of a healthy BM annotation reference. (A) UMAP obtained from clustering analysis of five healthy BM samples (7,161 cells) with expression-based annotations. HSC, hematopoietic stem cells; LMP, lympho-myeloid progenitors; MDP, monocyte/dendritic cell progenitors; NP, neutrophil progenitors; EBMP, eosinophil/basophil/mast cell progenitors; BCP, B cell progenitors; MEP/MKP, megakaryocyte/erythrocyte and megakaryocyte progenitors; EP-I and -II, erythrocyte progenitors; EP-Cy, cycling erythrocyte progenitors. (B-C) Dot plots of expression of (B) genes and (C) proteins used to annotate the UMAP in Figure 2A. Colors represent average expression and dot size percentage of cells in each cluster expressing the gene or protein. (D) Separate clustering analyses of CML diagnosis samples analyzed within the study (n=16; 1,250-5,554 cells per sample). Coloring and annotation based on cell label transfer from the healthy reference annotation in Figure 2A. (E) UMAPs from combined clustering analyses of diagnosis and TKI follow-up samples from the same patient. The figure illustrates the location of cells annotated as HSC by the healthy reference (top panel) as well as CD90 and CD38 expression across each patient-specific UMAP (mid and bottom panel, respectively). (F-J) Generation of a CML-specific annotation reference. (F) UMAP from combined clustering of the Figure 2E samples (18,078 CML diagnosis and 4,104 TKI follow-up cells). (G) Healthy reference cell label transfer annotation of the Figure 2F UMAP. (H) Feature plot visualization of protein expression of healthy and CML-specific stem cell markers. (I) Distribution of *BCR-ABL1*-expressing cells. (J) Final CML-specific annotation reference.

Next, we performed patient-specific clustering of cells from CML samples and used the healthy reference to annotate the CML patient cells by cell label transfer. Separate UMAP clustering visualizations of diagnosis HSPC from each CML patient are shown in Figure 2D. Notably, in seven patients, combined analysis of diagnosis and follow-up samples (Supplemental Figure 2D) revealed two separate clusters annotated as HSC with high and low expression of CD90 and CD38, respectively (Figure 2E). We speculated that these two clusters may represent separate HSC and CML LSC populations, and next aimed to set up a CML-specific annotation reference that distinguished these subpopulations within the CML stem cell compartment.

To this end, cells from the seven patients were combined, and a new clustering analysis was performed (Figure 2F). Also, the combined UMAP harbored two separate groups of cells annotated as HSC by the healthy reference (clusters 8 and 14; Figure 2G). While CML diagnosis cells appeared in both clusters, cells obtained during TKI treatment mostly resided in cluster 8 (Supplemental Figure 3A). Clusters 8 and 14 both showed expression patterns characteristic of HSC, including expression of CD90 and low/no expression of CD38 and CD45RA, but cluster 14 additionally showed expression of the previously reported CML LSC markers CD26^15^ and CD25^16^ (Figure 2H). In addition, cells identified as *BCR-ABL1*^+^ almost exclusively resided in cluster 14 (Figure 2I), further supporting that this cluster harbors the LSC. Clusters 8 and 14 were thus denoted HSC and LSC, respectively. Annotation of remaining clusters (Figure 2J) was performed as described for the healthy reference in the Supplemental Methods. Annotation marker feature plots and dot plots, as well as the cell cycle analysis used for annotation of the cycling clusters, are shown in Supplemental Figure 3B-F and Supplemental Tables 9-10.

### Myeloid bias of the HSPC compartment in CML diagnosis samples

Next, we clustered cells from CML diagnosis and follow-up samples together with those from healthy donors (70,414 cells total) and used the CML-specific reference for annotation (Figure 3A). The addition of cells from another nine CML patients and five healthy controls did not substantially alter the appearance relative to the Figure 2J UMAP, apart from the appearance of a B cell progenitor (BCP) cluster predominantly stemming from the healthy and follow-up BM samples (Supplemental Figure 4A). The expression patterns of traditional HSC and LSC markers (Figure 3B) as well as the location of *BCR-ABL1*^+^ cells (Figure 3C) further supported the existence of separate HSC and LSC populations.

**Figure 3.**
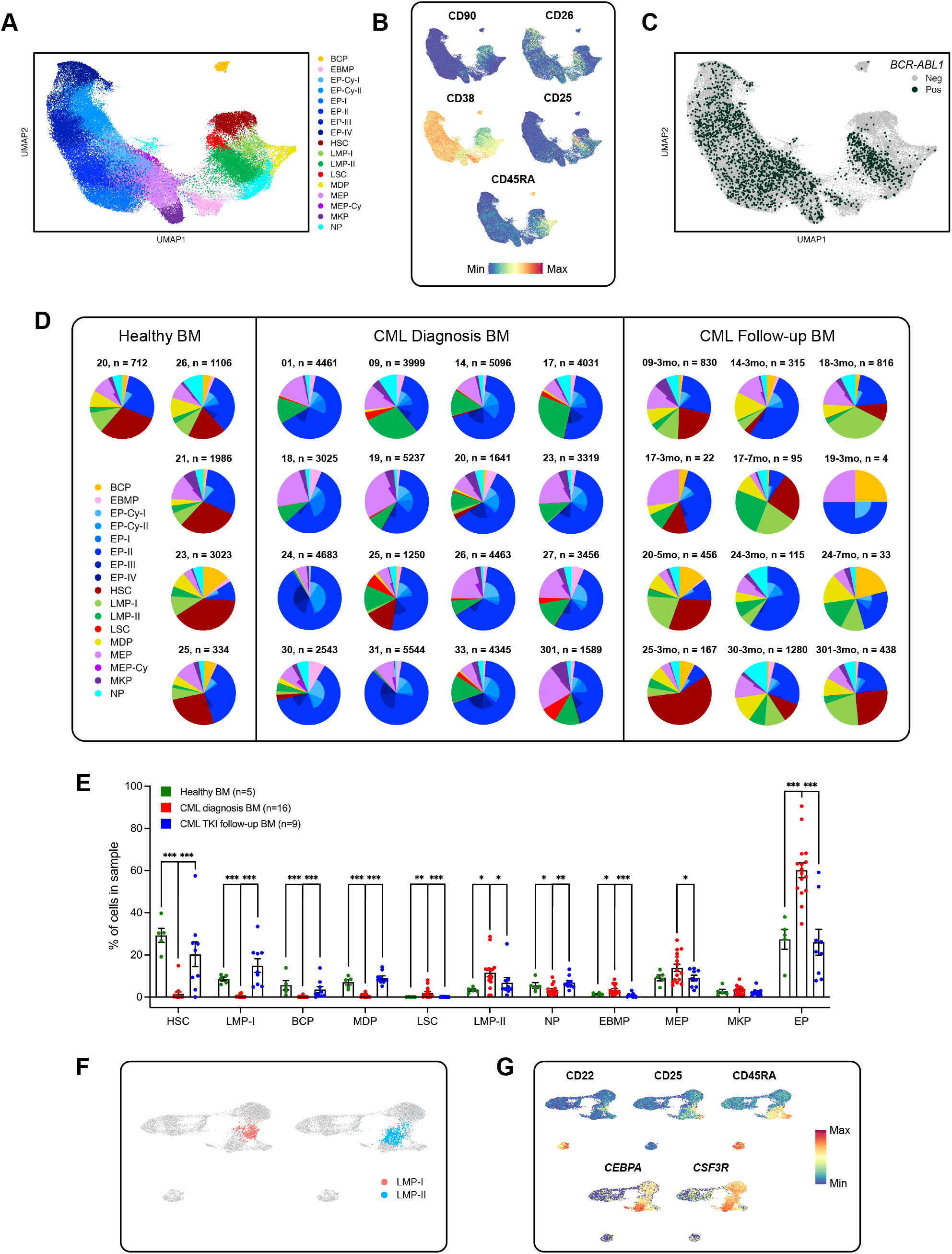
Multiomic analysis of the CD14^-^CD34^+^ stem and progenitor cell compartment at CML diagnosis and during TKI treatment. (A) UMAP visualization of combined clustering analysis of all healthy BM (5 samples; 7,161 cells), CML diagnosis (16 samples; 58,682 cells) and TKI follow-up samples (12 samples; 4,571 cells), annotated using cell label transfer from the ‘CML-specific annotation reference’. (B) Expression of healthy and CML-specific stem cell markers. (C) Distribution of *BCR-ABL1*-expressing cells. (D) CD14^-^CD34^+^ cell type proportions in each sample analyzed in the study. In the outer pie chart, erythrocyte progenitors are grouped and shown in blue and megakaryocyte/erythrocyte progenitors in lilac. (E) Quantification of cell type heterogeneity within the CD14^-^CD34^+^ BM compartment at CML diagnosis, during TKI treatment and in healthy controls. Each dot represents an individual sample. Samples with less than 50 cells were excluded from analysis. *p<0.05, **p<0.01, ***p<0.001. Error bars represent SEM. (F) Location of cells labeled as LMP-I and -II when using the CML-specific reference to annotate the healthy reference UMAP. (G) Lymphoid (CD22, CD25, CD45RA) and myeloid (*CEBPA, CSF3R*) gene/protein expression across the healthy reference UMAP.

To assess the composition of the CD14^-^CD34^+^ HSPC compartment at CML diagnosis, during TKI treatment and in healthy BM, we determined cell type proportions in each sample. As shown in Figure 3D-E, the CML diagnosis HSPC displayed rather homogeneous patterns of enriched cell types as shown in the left part of the Figure 2A UMAP, including LSC, LMP-II, erythrocyte progenitors (EP), and megakaryocyte/erythrocyte progenitors (MEP). Conversely, cell types on the right-hand side (*i.e*., HSC, LMP-I, monocyte/dendritic cell progenitors (MDP) and BCP) were almost absent in CML diagnosis BM. When using CML-specific reference-based cell label transfer to annotate healthy BM cells (Supplemental Figure 4B) LMP-I cells tended to aggregate on the right side of the healthy reference LMP cluster, while LMP-II cells were mostly found on the left side (Figure 3F). LMP-I displayed expression of CD22, CD25 and CD45RA (Figure 3G), suggesting a lymphoid bias. LMP-II instead showed myeloid bias, *e.g.* expressing *CEBPA* and *CSF3R* (Figure 3G) and connecting to the NP cluster. The myeloid bias of the LMP-II population was further supported by differential expression analysis of cells annotated as LMP-I and LMP-II in the healthy reference UMAP, where *CEBPA* and *CLEC12A* were among the top upregulated genes in LMP-II cells (Supplemental Table 11).

Almost all CML patients showed high proportions of erythrocyte progenitors at diagnosis (EP; Figure 3E). In addition, most patients harbored large numbers of the myeloid-biased LMP-II, whereas the lymphoid-biased LMP-I compartment was absent in the CML diagnosis BM. The few patients with LMP-I cells at diagnosis (CML20, 25 and 30) also harbored a population of HSC along with B and/or monocyte/DC progenitors (BCP and MDP; Figure 3D). As these populations were predominantly *BCR-ABL1*^-^ (Figure 3C), this may imply higher presence of normal hematopoiesis in this subgroup of patients. Samples obtained during TKI treatment more closely resembled the HSPC compartment in healthy BM with re-appearance of HSC and lymphoid progenitors.

We thus observed a strong myeloid bias in the diagnosis CML HSPC compartment with excessive proliferation, not only of granulocytic progenitors, but also of cells within the erythroid compartment. Interestingly, although the healthy UMAP in Figure 2A suggested early branching and generation of megakaryocytic/erythrocyte progenitors directly from the HSC population^17–19^, analysis of the CML diagnosis BM was compatible with a more traditional view of hematopoiesis, with all myeloid populations stemming from a common progenitor (here LMP-II). This aspect was particularly apparent when only CML diagnosis samples were included in the clustering analysis (Supplemental Figure 4C).

### The most immature CML LSC express VWF and TIM3

The most generally accepted LSC phenotype in CML is the expression of CD34 and lack of CD38^20^. To increase the resolution of stem cell analysis, we performed reclustering of the CD14^-^CD34^+^CD38^-/low^ cell compartment. CD34 and CD38 protein expression data, which are less flawed by drop-outs due to transcriptional bursting than transcriptomic data, were utilized for cell selection. Since the absolute signal in the protein data may vary between experiments, we performed separate gating of CD34^+^CD38^-/low^ cells processed on each cartridge (Figure 4A).

**Figure 4.**
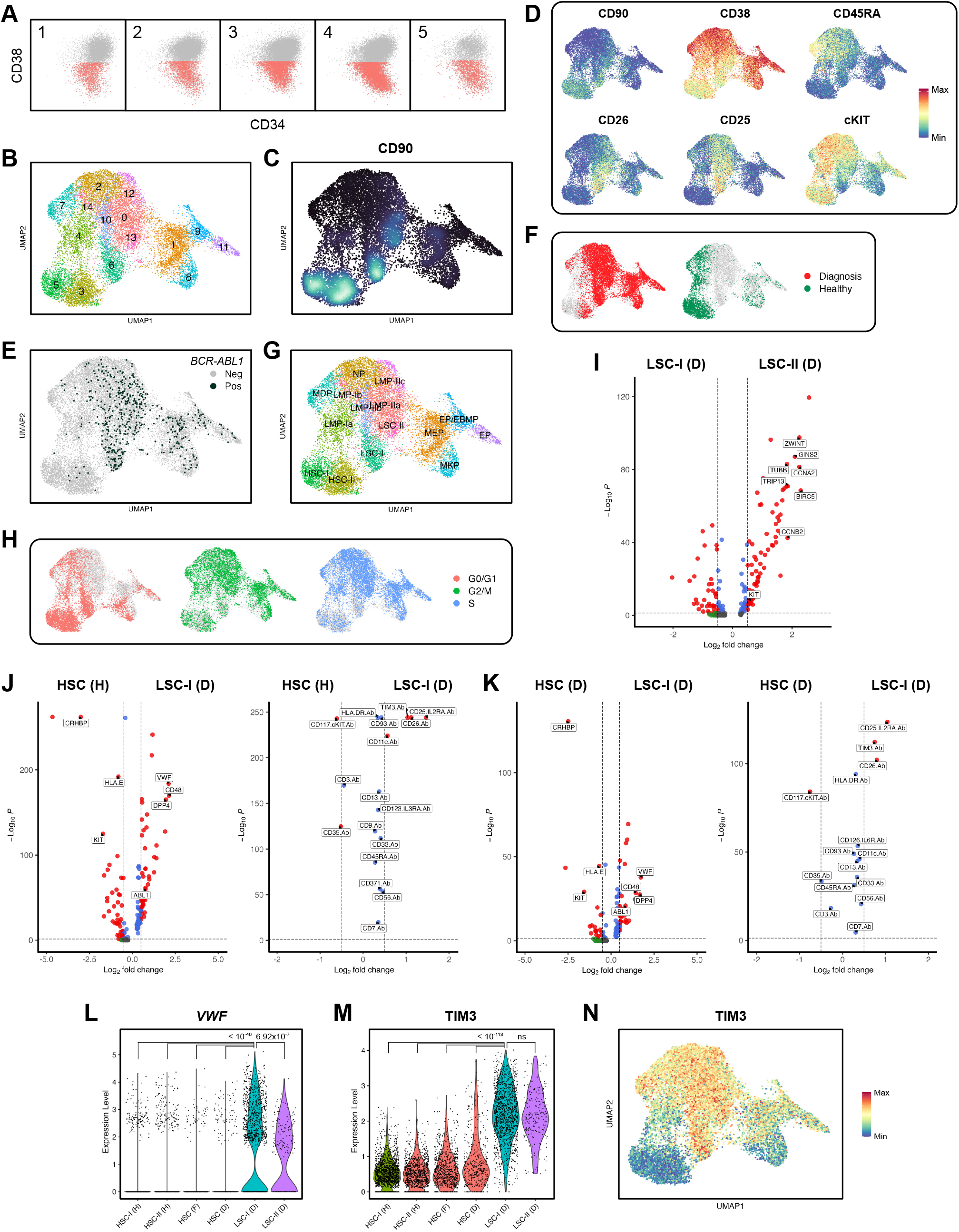
Multiomic analysis of the traditional CD34^+^CD38^-/low^ CML LSC/HSC compartment. (A) Cartridge-specific AbSeq-based gating for the analysis of CD34^+^CD38^-/low^ cells in CML patient and healthy BM. (B) UMAP of CD14^-^CD34^+^CD38^-/low^ cells from all healthy BM (5 samples; 4,090 cells), CML diagnosis (16 samples; 11,247 cells) and TKI follow-up samples (10 samples; 2,483 cells). (C) Density nebulosa plot showing the expression of CD90 across the CD14^-^CD34^+^CD38^-/low^ UMAP. (D) Feature plot visualization of the expression of healthy and CML-specific stem cell markers. (E) Distribution of *BCR-ABL1*-expressing cells across the CD14^-^CD34^+^CD38^-/low^ UMAP. (F) Location of cells deriving from CML diagnosis and healthy BM samples. (G) Annotation of the CD14^-^CD34^+^CD38^-/low^ UMAP. (H) Location of cells in different cell cycle phases according to Seurat’s CellCycleScoring function. (I) Volcano plot from a differential gene expression analysis comparing diagnosis LSC-I (LSC-I (D)) and -II (LSC-II (D)). Dashed lines indicate adjusted p values of 0.05 and log_2_FC=+/-0.5. (J-K) Volcano plots from differential gene (left panels) and protein (right panels) expression analyses comparing (J) healthy HSC (healthy BM cells in clusters HSC-I and -II; HSC (H)) to CML diagnosis LSC-I (LSC-I (D)), and (K) CML diagnosis HSC (diagnosis BM cells in clusters HSC-I and -II; HSC (D)) to CML diagnosis LSC-I (LSC-I (D)). Dashed lines indicate adjusted p values of 0.05 and log_2_FC=+/-0.5. (L-M) Violin plots showing (L) *VWF* and (M) TIM3 expression in stem cell populations from healthy (‘(H)’), CML diagnosis (‘(D)’) and follow-up (‘(F)’) BM samples. ‘HSC (F)’ and ‘HSC (D)’ comprise both HSC-I and HSC-II from the respective sample types. (N) Feature plot of TIM3 expression across the CD14^-^CD34^+^CD38^-/low^ UMAP.

The reclustering of the CD14^-^CD34^+^CD38^-/low^ cell compartment gave rise to 15 clusters (Figure 4B). The identities of the selected cells are highlighted in Supplemental Figure 5A with the annotation of the reclustered UMAP by the CML-specific reference shown in Supplemental Figure 5B. Notably, this UMAP comprised four distinct immature CD34^+^CD38^-/low^CD90^+^CD45RA^-^ cell populations (clusters 5, 3, 6 and 13; Figure 4B-D). Clusters 6 and 13 harbored *BCR-ABL1*^+^ cells expressing the LSC markers CD25 and CD26 (Figure 4D-E) suggesting leukemic origin, while clusters 5 and 3 appeared healthy. This notion was further supported by the location of cells from healthy donors versus CML diagnosis samples (Figure 4F). Annotation of the CD14^-^CD34^+^CD38^-/low^ UMAP (Figure 4G) was performed as previously described. Feature plots for genes and proteins used in the annotation are provided in Supplemental Figure 5C-D.

We next used Seurat’s CellCycleScoring function to test whether the two clusters of leukemic stem cells, provisionally termed LSC-I and LSC-II, differed in cell cycling status. In this analysis, cells in the LSC-I cluster were predominantly non-cycling (G0/G1) whereas LSC-II cells showed S/G2/M phase-related gene expression (Figure 4H). Furthermore, after removing outlier cells clustering far from the main HSC and LSC clusters (Supplemental Figure 5E), a differential expression analysis between type I and II LSC showed that seven out of ten of the top upregulated genes in LSC-II cells were cell cycle-related (Figure 4I, Supplemental Table 12-13). Quiescent LSC are reportedly more difficult to target with TKI therapy^7,8^. In agreement, the non-cycling LSC-I population displayed the previously reported TKI-resistance phenotype (CD34^+^CD38^-/low^CD45RA^-^cKIT^-^CD26^+^; Figure 4D)^8^. LSC-II cells, on the other hand, showed significantly higher expression of cKIT on the gene and protein level (Supplemental Table 12-13). These results thus point to the presence of two types of LSC at CML diagnosis, one in active proliferation and one more quiescent.

We next set out to compare gene and protein expression in the identified quiescent and TKI-resistant CML LSC (LSC-I) with expression in HSC from healthy controls (Figure 4J, Supplemental Tables 14-15) or HSC in CML BM at diagnosis (Figure 4K, Supplemental Tables 16-17). As expected, LSC-I cells displayed upregulation of previously reported LSC markers, such as CD25 and CD26. In addition, LSC-I had significantly lower levels of *CRHBP*, CD35 and cKIT when compared with HSC. Notably, *ABL1* expression was significantly higher in the LSC, likely reflecting *BCR-ABL1* transcripts incorrectly identified as *ABL1* transcripts in the standard CITE-seq workflow.

*VWF* was a top LSC-upregulated gene in these comparisons (Figure 4L). Other top differentially regulated genes included *HLA-E* and *CD48* that encode structures of relevance to NK cell recognition^21,22^. At the protein level, TIM3 was significantly upregulated in LSC compared with HSC from both healthy donors and TKI-treated patients (Figure 4M). A feature plot displaying TIM3 expression across the CD14^-^ CD34^+^CD38^-/low^ UMAP indicated preferential expression among LSC, LMP-II, NP and MDP (Figure 4N). While TIM3 has previously been reported to be upregulated in AML stem cells^23,24^, little is known regarding its role in CML.

### Previously reported CML LSC markers are surface expressed on the immature LSC-I population to different extents

We here report the first comprehensive transcriptional characterization of the CML CD14^-^CD34^+^ compartment with parallel surface protein expression data. In view of the observed transcriptome-based cell type heterogeneity within the traditionally defined CD34^+^CD38^-/low^ CML LSC compartment, we assessed the distribution of expression of previously reported CML LSC markers^20,25,26^ on the surface of transcriptionally defined LSC and HSC populations within the CD34^+^CD38^-/low^ compartment (Figure 5). While most of these markers indeed were expressed by the LSC-I population, CD26 showed the highest specificity for these cells within the CD34^+^CD38^-/low^ compartment, while the specificity was lower for CD9, CD25, CD33, CD93, CD123 and CD133 (Figure 5). Expression patterns for the other proteins in the panel across the CD14^-^CD34^+^CD38^-/low^ UMAP are shown in Supplemental Figure 6.

**Figure 5.**
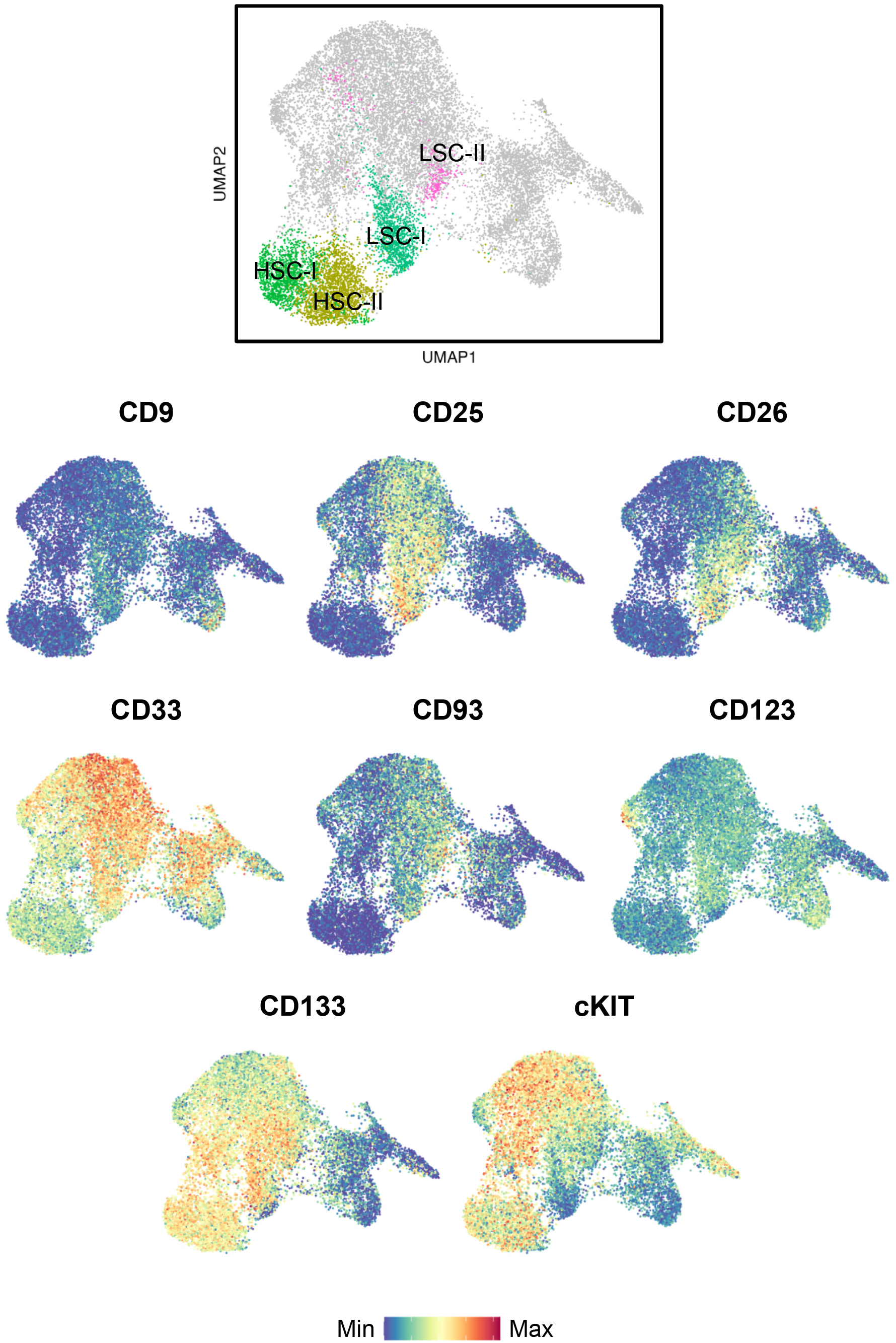
Expression of previously reported CML LSC markers on the surface of CD14^-^CD34^+^CD38^-/low^ cells from healthy and CML samples. The top panel illustrates the location of the four stem cell populations identified within the CD14^-^CD34^+^CD38^-/low^ compartment of these samples.

## Discussion

Despite the outstanding success of tyrosine kinase inhibitors in CML, most patients require life-long therapy and monitoring, which is associated with impaired quality of life and significant societal costs. Thus, strategies to target the TKI-resistant LSC population are needed to enable permanent cure. Recent years have seen the development of single-cell technologies that hold promise for in-depth characterization of the CML LSC compartment. CITE-seq captures gene and protein expression within the same cell and may enable identification of unique or overexpressed surface markers of transcriptionally defined populations, allowing subsequent isolation and functional characterization and/or identification of therapeutic targets. In CML, proteo-transcriptomic profiling of LSC should ideally be combined with detection of the disease-causing translocation *BCR-ABL1* for definitive distinction between leukemic and healthy cells.

Current high-throughput scRNAseq methods are by design not compatible with genotyping of aberrations far from the transcript ends such as the *BCR-ABL1* fusion point, and previous studies addressing transcriptomic heterogeneity among CML LSC have been limited by either lacking simultaneous protein expression analysis, *BCR-ABL1* detection or by a low number of cells and/or markers^7–9^. We here present a method allowing detection of the *BCR-ABL1* fusion point in cells with paired CITE-seq data, thus enabling high-throughput proteo-transcriptomic profiling of leukemic cells. The *BCR-ABL1* transcript detection sensitivity largely corresponded to that of the previously described circularization GoT method^10^, and although the detection level was relatively low, this strategy satisfactorily allowed distinction between clusters of HSC and CML LSC.

The obtained multiomic CITE-seq dataset, comprising >70 000 primary HSPC, allowed in-depth analysis of the CD34^+^CD38^-/low^ compartment in CML. While the Lin^-^CD34^+^CD38^-/low^ population is frequently regarded as one compartment of HSC/LSC in the literature, our study shows that the CD34^+^CD38^-/low^ compartment holds a multitude of cell types of varying lineage and maturity. Bulk HSC/LSC comparisons across sample types thus become challenging to interpret, highlighting the advantage of current single-cell approaches. When zooming in on the most immature populations among CD34^+^CD38^-/low^ cells, the CITE-seq data indicated presence of two distinct immature leukemic populations in CML (denoted LSC-I and LSC-II), both harboring *BCR-ABL1^+^* cells with expression of the LSC markers CD25 and CD26. However, only the LSC-I population displayed the CD34^+^CD38^-/low^CD45RA^-^cKIT^-^CD26^+^ phenotype that has previously been associated with LSC that persist during TKI treatment^8^. The quiescent LSC-I population is likely the population to target to enable permanent cure, and we thus compared its gene and protein expression to the distinct HSC populations. Analyses of surface expression of previously reported CML LSC markers (CD9, CD25, CD26 CD33, CD93, CD123, CD133 and cKIT^20,25,26^) indicated CD26 as the marker with the highest specificity and selectivity for LSC-I.

In differential expression analysis, a previously unreported elevated expression of *VWF* was detected in LSC-I. Since expression of *VWF* has been reported to mark a quiescent group of HSC with myeloid and platelet differentiation bias^27^, we speculate that its expression may be connected to the skewed myeloid proliferation that characterizes CML^8,28^. It remains, however, to be established whether *VWF* has any functional role for this phenotype.

We also identified significant cell surface overexpression of TIM3 when comparing CML LSC with normal HSC, which is in line with a previous report showing upregulated TIM3 gene expression (*HAVCR2*) in CML LSC based on mining of publicly available RNA-seq datasets^29^. In addition, TIM3 is reportedly overexpressed and critical for self-renewal of LSC in acute myeloid leukemia (AML)^23,24,30^. Collectively, these results point to the targeting of TIM3 as a possible strategy to reduce stem cell renewal in CML. A potential caveat is that TIM3 has been identified as a marker of myeloid differentiation commitment in healthy hematopoietic cells^31^. However, the most immature HSC population hardly expresses TIM3 and studies employing TIM3-targeting opsonizing antibodies or CAR T constructs to enhance elimination of leukemic stem cells in AML suggest that this indeed may be a viable approach^32,33^.

We observed that the CML CD14^-^CD34^+^ compartment at diagnosis was characterized by pronounced expansion of myeloid-biased LMP-II and erythrocyte progenitors with few immature cells and lymphoid progenitors. The enhanced presence of erythrocyte progenitors in CML BM is in line with previous reports^9,28^, but the high number of CD34^+^ cells analyzed and the presence of CITE-seq data additionally allowed assessment of differentiation patterns within the CML BM. These results are thus to our knowledge the first to suggest altered differentiation trajectories in the leukemic versus healthy BM. According to the classical model of hematopoiesis, differentiation of HSC into mature blood cells occurs in a stepwise fashion with the first bifurcation between common myeloid progenitors (CMP) and common lymphoid progenitors (CLP)^34^. However, recent reports suggest that megakaryocyte/erythrocyte progenitors (MEP) arise directly from HSC^17–19^. Accordingly, we observed transcriptional signs of direct differentiation from HSC to MEP in healthy BM. By contrast, the generation of MEP in CML instead seemed to occur through a myeloid-biased progenitor population. Hence, although conclusions from our healthy BM data cohere with those of other recent healthy BM scRNA-seq studies, the analysis of CML data indicates that hematopoiesis may take different routes depending on BM conditions.

In line with a previous report^8^, the collection of follow-up sample HSPC after TKI therapy indicated resemblance to healthy BM with a decreased myeloid output along with increased proportions of immature HSC and lymphoid-lineage progenitors. As most patients had attained complete cytogenetic responses (CCyR) at the point of follow-up sampling, we were unable to assess the LSC compartment during TKI treatment. The follow-up samples for patients who had not achieved CCyR contained low numbers of CD14^-^ CD34^+^ cells, preventing detailed analysis. An aim for future studies is thus collection of patient samples closer to the initiation of TKI treatment to potentially capture semi-refractory LSC.

In conclusion, we here provide an unprecedentedly detailed proteo-transcriptomic characterization of the LSC compartment in CML. By coupling *BCR-ABL1* expression detection to single-cell gene and protein expression data, we identified quiescent CML LSC displaying previously unreported elevated expression of TIM3 and *VWF*. Our results suggest that TIM3, a surface expressed antigen, may be targetable for eradication of TKI-eluding LSC.

## Acknowledgments

We gratefully acknowledge CML patients, healthy donors and medical professionals who allowed sample acquisition for this study. We would also like to thank past and present members of the BD Multiomics Alliance and the BD R&D department for collaborative efforts on the development of the multiomic panel and input on the optimization of the *BCR-ABL1* detection method. We are grateful to Gustav Holmgren at TATAA Biocenter for running the BluePippin PCR product isolation.

This work was supported by grants from the Swedish Research Council (2020-1437 (A.M.) and 2020-02783 (F.B.T.)), the Swedish Cancer Society (2018/582 (K.H.), 22 2128 Pj (A.M.) and 19 0449 Pj (F.B.T.)), the Swedish state via the ALF agreement (ALFGBG-718421 (K.H.), ALFGBG-724881 (A.M.), and ALFGBG-963642 (F.B.T)), the Assar Gabrielsson Foundation, the Wilhelm and Martina Lundgren Research Foundation and the Sahlgrenska Academy at the University of Gothenburg.

## Authorship

### Contribution

M.S.N., H.K. and J.G. designed, performed, and analyzed experiments; M.S.N. set up, optimized and performed experiments related to the *BCR-ABL1* detection method, with assistance from Z.S and H.K; M.S.N. designed and performed the bioinformatic analysis; O.R. and L.W. provided healthy and CML patient samples, respectively; H.K., M.S.N., Z.S., and S.P. isolated mononuclear cells from patient samples; F.B.T., A.M., L.W. and K.H. conceived, designed and supervised the study; M.S.N., K.H., A.M., H.K. and F.B.T. wrote the manuscript, and all authors edited the manuscript.

