## Supplemental methods for "Multiomic single-cell analysis identifies von Willebrand factor and TIM3-expressing *BCR-ABL1*^+^ CML stem cells"

*Cell lines*

The blast crisis CML cell line K562 was cultured in Iscove's Modified Dulbecco's Medium (Gibco) supplemented with 10% heat inactivated fetal calf serum (FCS), 1% L-glutamine, 1% sodium pyruvate and 1% PEST. Cells were cryopreserved in aliquots and stored at -140 to -180°C for use of transcriptionally similar cells in different experiments.

*Samples and isolation of bone marrow mononuclear cells*

The study included a total of 16 chronic phase CML patients and five healthy donors. CML patient bone marrow (BM) samples were aspirated from the posterior iliac crest at diagnosis (n=16) and three to seven months into TKI treatment ('follow-up samples'; twelve samples obtained from ten patients) at the Sahlgrenska University Hospital (Gothenburg, Sweden) and Uddevalla Hospital (Uddevalla, Sweden). Healthy BM samples were obtained from the femur of osteoarthritic patients undergoing hip replacement surgery at the Sahlgrenska University Hospital (Mölndal, Sweden). Exclusion criteria included previous chemotherapy-treated cancer and ongoing systemic treatment with steroids. Clinical details of the CML patients, as well as age and sex of healthy donors, are provided in Supplemental Table 1. The study was approved by the Regional Ethics Committee in Gothenburg (ethical application numbers: 011-17 and T1104-18) and all patients gave written informed consent to participation in the study in accordance with the Declaration of Helsinki.

BM samples were diluted in buffered saline, layered on Lymphoprep (STEMCELL Technologies) and BM mononuclear cells (MNC) were isolated by density gradient centrifugation. Removal of co-sedimenting immature $CD15^+$ granulocytes and/or enrichment of $CD34^+$ cells was accomplished by morphological MNC FACS sorting, negative selection of CD15-expressing and/or positive selection of CD34-expressing cells using magnetic microbeads (Miltenyi Biotec). Cells were cryopreserved and kept at -140 to -180°C until use.

*Single-cell gene and protein expression analysis and panel design*

The BD Rhapsody Single-Cell Analysis System (BD Biosciences) was used for parallel targeted single-cell CITE-seq analysis of 597 genes and 51 surface proteins (Figure 1A). The panels (Supplemental Tables 2-4) were developed in collaboration with the labs of prof. M. Rieger (Goethe University Hospital, Frankfurt, Germany) and Dr. B. Paiva (University of Navarra, Pamplona, Spain).

*Sample preparation for multiomic expression profiling*

Samples were processed in five batches on five BD Rhapsody cartridges. Importantly, each cartridge had one healthy control sample, diagnosis samples were spread across all five cartridges and CML follow-up samples were always added to the same cartridge as the corresponding diagnosis sample to minimize potential batch effects on multiomic expression results.

Cryopreserved BM samples were thawed in a 37°C water bath and dropwise mixed with RPMI-1640 (Gibco) with 10% heat inactivated (hi) FCS. Samples were centrifuged at 350 g, 7 min, 4°C and washed once with RPMI-1640, 10% hi FCS. Cells were resuspended in PBS with 0.5% BSA and 0.1% EDTA and filtered through 35 µm cell strainers. After Sysmex XP-300 (Sysmex) cell counting, a maximum of 10 M cells per sample were used in subsequent steps. After another 350 g, 7 min centrifugation, samples were resuspended in 180 µl BD Stain Buffer (FBS)(BD Biosciences) and stained with 20 µl sample-specific oligo-conjugated cell hashing antibodies (‘Sample Tags’; BD Single-Cell Multiplexing kit, BD Biosciences) for 20 min at room temperature (RT) and washed twice with BD Stain Buffer (FBS).

Samples that had not been CD34 purified prior to cryopreservation were subjected to positive $CD34^+$ cell selection using magnetic microbeads, LS columns and MidiMACS separators (Miltenyi Biotec) using the manufacturer’s instructions (one fourth of the recommended kit reagent volumes for 100 M cells). After Bürker chamber cell counting, $CD34^+$ cells from the different samples were pooled, aiming to achieve 1:1 ratio of individual CML diagnosis as well as healthy BM samples. For the CML follow-up samples, all $CD34^+$ cells were typically added due to lower sample cell numbers.

The cell pool was subjected to cell surface co-labeling with fluorescent, unconjugated and BD AbSeq oligo-conjugated antibodies (BD Biosciences). The lists of antibodies used are provided in Supplemental Tables 4 and 5. Staining was performed in BD Stain Buffer (FBS) at a total staining volume of 200 µl, using 2 µl of each BD AbSeq antibody and fluorescent/unconjugated antibodies at a 1:1 molar ratio with the corresponding AbSeqs. Of note, CD14, CD34 and CD38 fluorescent antibodies were used for subsequent FACS sorting and analysis, whereas the unconjugated HLA-ABC antibody was used to mute the signal from the HLA-ABC AbSeq antibody, which otherwise tends to dominate downstream single-cell Abseq signals. Cells were incubated with the antibodies for 45 minutes on ice, washed twice with BD Stain Buffer (FBS) and resuspended in PBS with 0.5% BSA and 0.1% EDTA in preparation for cell sorting.

To remove cell debris, singlet CD14$^-$CD34$^+$ BM MNC were sorted into collection tubes containing cold RPMI-1640 with 10% hi FCS using a BD FACSAria Fusion cell sorter (BD Biosciences) in yield mode. After sorting, cells were centrifuged at 400 g, 5 min, 4°C, resuspended in 198 or 620 µl cold Sample Buffer from the BD Rhapsody Cartridge Reagent Kit (BD Biosciences) and viability stained with Calcein AM (Thermo Fisher Scientific) and Draq7 (BD Biosciences) according to the instructions in the *Single-Cell Capture and cDNA Synthesis with the BD Rhapsody Single-Cell Analysis System* protocol (Doc ID: 210966, rev 1.0; BD Biosciences).

Single-cell capture was performed on the BD Rhapsody Single-Cell Analysis System in accordance with the above-mentioned protocol, aiming to capture a total of 45,000 cells per cartridge. At cell load to the BD Rhapsody Express System, the viability of the samples was consistently above 92% (92.2-97.5%). On four out of five cartridges, 5,000 cells of the *BCR-ABL1*$^+$ K562 cell line were included as positive controls for *BCR-ABL1* transcript detection. These cells were thawed and processed in parallel with the patient BM samples, using a similar approach.

*Sequencing library preparation for multiomic expression profiling*

Single-cell capture beads carrying mRNA, BD AbSeq and BD Sample Tag oligos were pooled for reverse transcription, exonuclease treatment and subsequent parallel preparation of mRNA, BD AbSeq and Sample Tag Illumina sequencing libraries.

Reverse transcription was performed in 100 µl reactions using 1000 U SuperScript IV Reverse Transcriptase (ThermoFisher Scientific), 200 U Ribonuclease inhibitor (ThermoFisher Scientific), 1x SSIV buffer (ThermoFisher Scientific), 0.5 mM dNTP mix (ThermoFisher Scientific), 5 mM DTT (ThermoFisher Scientific) and 6 µl Bead RT/PCR Enhancer (BD Biosciences; BD Rhapsody cDNA Kit). The bead suspension was incubated on a ThermoMixer C instrument with a SmartBlock Thermoblock 1.5 mL (Eppendorf) at 1,200 rpm and 50°C for 50 minutes. Exonuclease treatment was performed according to BD's protocol using the reagents in the BD Rhapsody cDNA kit (BD Biosciences). After cDNA synthesis and exonuclease treatment, beads were stored in Bead Resuspension Buffer (BD Rhapsody cDNA kit) at 4°C until sequencing library preparation.

mRNA targeted, BD AbSeq and Sample Tag sequencing libraries were prepared using the BD Rhapsody Targeted mRNA and AbSeq Amplification Kit following the manufacturer's instructions (Doc ID: 214508 Rev 3.0; BD Biosciences). Sequences of the primers used can be found in Supplemental Tables 2-3. Cartridge-specific reverse primers were used in the index PCR, allowing for pooling and sequencing of all libraries in the same sequencing run.

Prior to sequencing, libraries were quality controlled on a 4200 TapeStation System (Agilent) using High Sensitivity D1000 ScreenTape and Reagents (Agilent) to ensure correct size and purity. The Qubit dsDNA HS Assay kit (ThermoFisher Scientific) was used to assess library concentrations. Libraries were pooled aiming to achieve a sequencing depth of 10,000 reads per cell for mRNA libraries, 2,000 reads per cell for sample tag libraries and 500 reads per cell per antibody for AbSeq libraries.

*BCR-ABL1 sequencing library preparation*

To assess *BCR-ABL1* expression in the single cells analyzed with the BD Rhapsody Single-Cell Analysis System, we developed a method similar to that reported by Nam et al ('Circularization GoT')[1]. The sequences of the primers used in our adaptation of the workflow are provided in Supplemental Table 6. Of note, primers were designed to allow detection of both major *BCR-ABL1* transcript variants (*i.e.* e13a2 and e14a2), only differing in the size of the final library fragment generated (with the e13a2 fragment being 75 bp shorter).

Following completion of PCR1 in the BD Rhapsody library preparation workflow and subsequent removal of the majority of the BD Rhapsody PCR1 products, the transcript-carrying beads were reused for *BCR-ABL1* sequencing library preparation (Figure 1B). In a first PCR (PCR-A), the *BCR-ABL1* cDNA on the beads from each cartridge were amplified in four 50 µl reactions using a primer binding upstream of the fusion point. Each 50 µl reaction contained 1x KAPA HiFi HotStart ReadyMix (Roche Diagnostics), 3 µl Bead RT/PCR Enhancer (BD Biosciences; part of the BD Rhapsody Targeted mRNA and AbSeq Amplification Kit), BCR-binding primer (IDT) at 0.5 µM and Universal Oligo (BD Biosciences) at 1 µM. Thermal cycling was performed using a Veriti 96-well fast thermal cycler (Applied Biosystems), with the following protocol: 95°C for 3 min, 27 cycles of 98°C for 20 s, 60°C for 30 s and 72°C for 4 min, and 72°C for 6 min. Notably, beads were pre-mixed immediately before placement in the thermocycler at 95°C in agreement with BD's protocol for the BD Rhapsody bead-based PCR1 (Doc ID: 214508, rev 3.0; BD Biosciences). After PCR completion, products from the four cartridge-specific PCR-A reactions were pooled, and short unspecific products were removed using Agencourt AMPure XP beads (Beckman Coulter) at a 0.7X ratio. To remove remaining longer unspecific products, samples were subjected to BluePippin (Sage Science) electrophoretic separation and collection of 5,950 bp products in 'tight' collection mode at TATAA Biocenter (Gothenburg, Sweden). The 5,950 bp target was based on the size of the specific product (pre separation) when analyzed on a BioAnalyzer instrument (Agilent).

In the next PCR (PCR-B), complementary overhangs (sequences from [1]) were added to the *BCR-ABL1* fragments in 50 µl reactions containing 1x KAPA HiFi HotStart ReadyMix and 0.3 µM forward and reverse primer (IDT). The thermal cycling protocol was 95°C for 3 min, followed by 6-7 cycles of 98°C for 20 s, 65°c for 30 s, and 72°C for 4 min, and 72°C for 6 min. Unspecific products were removed using 0.6X AMPure XP beads.

PCR-B products were circularized in 1,000 µl reactions containing 1X CutSmart Buffer (New England BioLabs), 10 µl NEBuilder Hifi DNA Assembly Master Mix (New England BioLabs) and 100 ng specific PCR-B product (based on Qubit concentration and BioAnalyzer purity). Samples were incubated at 50°C for 60 minutes in a Grant Bio PCH-1 heat block (Grant Instruments). After cooling on ice, 6U NEB Lambda Exonuclease (New England BioLabs) was added and samples were incubated at 37°C for 30 min, followed by enzyme inactivation at 65°C for 20 min.

Circularized PCR-B products were concentrated by ethanol (EtOH) precipitation. In brief, 40 µl 3M Na-Acetate pH 5.2 (4°C) and 1,000 µl 100% EtOH (-20°C) was added to 400 µl PCR product. Samples were incubated at -20°C for 1 h, followed by recovery of precipitated cDNA by centrifugation at 16,600 g, 30 min, 4°C. After washing with -20°C 70% EtOH (16,600 g, 10 min, 4°C), the supernatant was removed, and precipitated cDNA air-dried for approximately 15 mins. Pellets were resuspended in TE buffer (ThermoFisher Scientific) and placed at 4°C until PCR-C.

In the next PCR (PCR-C), the majority of the *ABL1* portion of the transcript was removed from the circularized products using a new set of complementary overhang primers. The reaction components and concentrations were the same as in PCR-B, but with another primer set. Thermal cycling was performed at 95°C for 3 min, 13-17 cycles of 98°C for 20 s, 65°C for 30 s and 72°C for 30 s, and 72°C for 1 min. PCR-C products were purified using AMPure XP beads at a 1.3X ratio.

A second circularization reaction was performed in the same way as the first but using 50 ng specific PCR-C product. After another round of exonuclease treatment and EtOH precipitation, PCR-D was used to linearize and create the final, shortened fragment for sequencing. PCR-D was performed the same way as PCR-B and -C, but with the following thermal cycling profile: 95°C for 3 min, 9 cycles of 98°C for 20 s, 65°C for 30 s and 72°C for 20 s, followed by 72°C for 1 min. PCR-D products were purified using AMPure XP beads at a 0.8X ratio.

In a final PCR (PCR-E), Illumina sequencing adapters and indices were added to the *BCR-ABL1* fragments using the Index PCR workflow and reagents in the BD Rhapsody library preparation protocol (Doc ID: 214508 Rev 3.0; BD Biosciences).

Following completion of each PCR as well as after BluePippin isolation, product lengths were verified using the Agilent High Sensitivity DNA kit on a BioAnalyzer instrument (Agilent) or the High Sensitivity D1000 ScreenTape and Reagents on a 4200 TapeStation System (Agilent).

*Sequencing library preparation for assessment of reverse transcription change*

To ensure that the change of reverse transcription procedure required for successful *BCR-ABL1* expression detection did not affect the performance of the general BD Rhapsody gene expression analysis, two K562 sequencing libraries were prepared in parallel using the original BD Rhapsody reverse transcription procedure ('BD RT', following the manufacturer's instructions) or the modified SuperScript IV reverse transcription procedure ('SSIV RT', as described under 'Sequencing library preparation for multiomic expression profiling'). Single-cell capture of K562 cells was performed on a single BD Rhapsody cartridge as previously described. The cell capture beads were split in two and subjected to the respective RT procedures. Following RT completion, exonuclease treatment and sequencing library preparation for the two libraries were performed in parallel according to previously mentioned protocols.

*Sequencing*

Sequencing of multiomic single-cell expression libraries was performed by the SNP&SEQ Technology Platform (Uppsala, Sweden; part of the National Genomics Infrastructure (NGI) Sweden and Science for Life Laboratory) using all four lanes of a S4-200 v1.5 flow cell on a NovaSeq 6000 instrument (Illumina) and a custom read setup of 64-8-0-74 (R1-i7-i5-R2). 20% PhiX control v3 (Illumina) was added to increase library complexity.

The paired single-cell *BCR-ABL1* detection libraries were sequenced on a MiniSeq instrument (Illumina) in-house (Sahlgrenska Center for Cancer Research, Gothenburg, Sweden), using the MiniSeq High Output Reagent kit (300-cycles; Illumina), a read setup of 151-8-0-151 and 35% PhiX control v3.

Sequencing of the RT assessment libraries was performed by NGI Stockholm (Stockholm; Sweden; part of the National Genomics Infrastructure (NGI) Sweden and Science for Life Laboratory) using the MiSeq Reagent Kit v3 (150-cycles; Illumina), a custom read setup of 64-8-0-74 (R1-i7-i5-R2) and 20% PhiX control v3 (Illumina).

*Data analysis and visualization*

Single-cell multiomic and *BCR-ABL1* detection fastq files from each cartridge were processed using the BD Rhapsody Targeted Analysis Pipeline (v. 1.10.1 and 1.10, respectively) on the Seven Bridges Genomics platform (https://www.sevenbridges.com). In short, the pipeline performs read quality filtering, read alignment, read pair combination and collapsing to raw molecules, RSEC (Recursive Substitution Error Correction) UMI adjustment, cell label filtering, sample of origin and multiplet determination (by sample tag reads) and generates single-cell gene and protein expression matrices. Within the pipeline, reads were aligned to reference files containing the final library fragment sequences of the targeted gene and AbSeq panels. Importantly, the target sequences for *BCR-ABL1* read alignment included only the portions closest to the *BCR-ABL1* fusion point, forcing mapping across the fusion to minimize false positive detection. Both sequencing datasets had >95% reads remaining after read quality filtering, and sequencing saturations for targeted mRNA, AbSeq and *BCR-ABL1* expression data exceeded 91% (91.6-98.3%) across all cartridges.

Following pipeline processing, the output RSEC UMI count files for multiomic and *BCR-ABL1* analysis for each cartridge were merged based on cell indices. Cells showing expression of at least one *BCR-ABL1* transcript were designated as *BCR-ABL1* positive in the following bioinformatic analysis.

Multiomic data analysis was performed in Rstudio (v. 2022.07.1+554; R v. 4.2.1) using the Seurat package (v. 4.2.0)[2]. In brief, cells were filtered based on genes expressed vs library size using SeqGeq software (BD Biosciences, v. 1.8.0)(Supplemental Figure 1). Gene and protein UMI count matrices for filtered cells were imported into Rstudio and used to create multiomic Seurat objects with mRNA and AbSeq assays. After quality control of gene contributions to the total UMI count per cell, cell cycle heterogeneity (Seurat's CellCycleScoring function using applicable 'cc.genes') and mitochondrial and ribosomal expression contributions to total UMI counts (consistently below 2 and 1%, respectively), confounding differences between proliferating cells (i.e. G2/M vs S phase) were regressed out of the data. mRNA expression data were normalized on a per cartridge basis using SCTransform[3], followed by merging (where applicable). The SCT assay was used for principal component analysis using variable genes selected by the sctransform function. Suitable numbers of principal components for graph-based clustering were assessed using the ElbowPlot function. The selected number of principal components (23-26) were used to construct a K-nearest neighbor (KNN) graph, followed by Louvain algorithm-based clustering. Finally, visualization of the clusters was accomplished using Uniform Manifold Approximation and Projection (UMAP).

Log-normalized RNA and centered log ratio (CLR) normalized AbSeq counts were used for differential expression analysis and visualization. Cluster annotations were manually determined based on differentially expressed genes and proteins using the FindAllMarkers function and visually by FeaturePlot expression patterns across the UMAPs. Cell label transfer from healthy or CML-specific annotation references to query datasets was performed using the FindTransferAnchors and TransferData functions in Seurat. Selection of cells based on expression of certain proteins or location on the UMAP was performed using Seurat's CellSelector function. Visualizations were generated in R using the ggplot2 (v. 3.4.0), SCPubr (v. 1.0.1 )[4], EnhancedVolcano (v. 1.14.0)[5], and RColorBrewer (v. 1.1.3)[6] packages, and in GraphPad Prism (v. 9.4.1).

*Generation of annotation references*

The clusters in the healthy annotation reference in Figure 2A were annotated based on previous publications[7–18]. As illustrated in Figure 2B-C, hematopoietic stem cells (HSC) were annotated based on the expression of CD90, *CRHBP*, *HLF*, and *MLLT3* and low expression of CD38 and CD45RA, lympho-myeloid progenitors (LMP) by decreased expression of HSC markers in parallel with increased expression of early myeloid/lymphoid markers such as *SPINK2* and *FLT3*, B cell progenitors (BCP) by their expression of CD10 (*MME*), CD19 and CD22, monocyte/dendritic cell progenitors (MDP) by expression of *KLF4*, *SAMHD1*, CD123 (*IL3RA)*, CD64, CD45RA and CD371, neutrophil progenitors (NP) by their expression of *CSF3R*, *CEBPA, CEBPE,* CD371 and cKIT, eosinophil/basophil/mast cell progenitors (EBMP) by expression of *HDC, GATA2* and *CEBPE* without concurrent expression of CD45RA, megakaryocyte/erythrocyte progenitors and megakaryocyte progenitors (MEP/MKP) by expression of *GATA2, MEIS1*, *MPL*, *VWF* and *GATA1,* and erythrocyte progenitors (EP) by expression of CD35, *GATA1, HBB,* and *AHSP*. Cycling EP (EP-Cy) were annotated based on showing similar expression patterns as EP-I, but with co-expression of cycling genes such as *POLA1*, *GINS2* and *RRM1,* along with being assigned as being in S/G2/M phase by Seurat's CellCycleScoring function in R (Supplemental Figure 2A).

The CML-specific annotation reference was generated in a similar manner. The validity of the annotation was assessed using the CML-specific reference to annotate the healthy cells in Figure 2A by cell label transfer (Supplemental Figure 4B). In this analysis, the CML-specific reference yielded annotations similar to the manual healthy reference annotations, supporting the robustness of the CML-specific annotations.

**References – Supplemental Material**
